## Supplementary information for "STRAINS: A Big Data Method for Classifying Cellular Response to Stimuli at the Tissue Scale"

Jingyang Zheng, Thomas Wyse Jackson, Lisa A. Fortier, Lawrence J. Bonassar, Michelle L. Delco, and Itai Cohen
*Cornell University*
(Dated: June 24, 2022)

### I. STRAIN-DEPENDENT CELLULAR RESPONSE

The strain field resulting from impact and the associated cell response had complex behaviors that varied spatiotemporally. For calcium, in the milliseconds after impact trauma (Green in Fig. 2a), increased concentrations can be seen in the cells proximal to the impact site. By one second, we observed spatially sporadic increases in calcium deeper in the imaged region. After a few seconds, the cells damaged at the impact site begin to fade, suggestive of death, while cells in the deeper regions exhibited a more uniform increase in calcium. Subsequently, the calcium concentration in cells below the impact slowly decayed. These differing calcium responses in cells that experienced various amounts of compression suggest that distinct mechanotransduction pathways may have been activated depending on mechanics governing the local strain field and the phenotype of each cell, which is known to vary with depth in cartilage [1–3].

Interestingly, in addition to this depth dependent response, we observed a different temporal pattern of calcium concentration in the region laterally adjacent to the impact (region indicated by orange boundary). Cells in this region near the articular surface primarily experienced shear strain during impact and showed increases in calcium on the tens of seconds time scale, with maximum intensities that were almost an order of magnitude lower. These data suggest that shear and compressive strains may trigger different mechanotransduction pathways, consistent with findings in cell-agarose constructs [4].

Such complex spatiotemporal patterns are also exhibited on longer time scales in all three measured channels (Fig. 2b): mitochondrial polarity (Red), calcium concentration (Green), and nuclear membrane permeability (Blue). We found that mitochondrial polarity rapidly diminished at the impact site in the minutes after injury. In the surrounding regions, we observed a slow decay in mitochondrial polarity over the course of multiple hours. Calcium concentration largely followed the same pattern, with some cells which exhibited transients on the scale of minutes (See supplementary Video 2). Conversely, nuclear membrane permeability initially showed a very low intensity throughout the region and reached higher intensities in a fraction of the cells in regions extending down to 400µm below the impact site on a time scale of hours. Finally, consistent with the short time calcium response, this pattern of cell death did not extend to areas of the tissue which experienced primarily shear strains.

Collectively, these distinct spatiotemporal patterns of cell response indicated that multiple mechanobiological pathways may have been activated in response to local strain. Cells at the impact site most likely died immediately due to membrane rupture caused by extremely high strains [5]. Cells further away from impact, which are subjected to lower strain intensities, may have activated different signaling pathways such as physiologic calcium signaling which led to normal tissue response (i.e. TRPV4 pathway [6, 7]) or superphysiologic calcium signaling which led to apoptosis (i.e. Piezo 1/2 [8–10]) each of which produced cellular signatures with characteristic combinations of fluorescence intensity curves. Developing an understanding of how such processes are related requires identifying distinct cellular signatures and mapping out where in the tissue they are localized. To obtain these maps, however, we must first identify each cell, track its movement and multi-channel fluorescence response over time (Fig.1e), and classify its cellular signature (Fig.1f).

#### A. Manual sorting of fluorescent intensity analysis in impacted articular cartilage shows spatially distinct cell behaviors

Below the impact site, we observed that many chondrocytes showed a high level or a rapid increase in their nuclear membrane permeability and low mitochondrial polarity, which most likely indicated that they were undergoing cell death. Interestingly, we observed different shapes in this channel. In some cells, nuclear membrane permeability was immediately elevated following impact and stayed high (Fig.4a). These cells were primarily localized in the 150µm just below the impact site. A second group showed a rapid increase in permeability in the 20 min following impact and maintained this intensity throughout the observation window (Fig.4b). These cells were distributed in a region extending 400 µm below the impact site. A third group showed a similar rapid increase in nuclear membrane permeability in the first 20 min following impact but then decayed in this channel to an intermediate level still higher than baseline. These cells could be found up to 1 mm below the impact site (Fig.4c). Additionally, in some cells down to 450µm below the impact site we observed a late rise in nuclear membrane permeability with no obvious trigger (Fig.4d), while in others we observed a rapid rise in nuclear membrane permeability despite the cell maintaining high mitochondrial polarity (Fig.4e).

There were a number of distinct behaviors associated with changes in cell calcium concentration. In the region located 150 - 400µm below the impact site, one group of cells showed a rapid drop in calcium concentration followed by a rapid increase in nuclear membrane permeability (Fig.4f). Another group showed calcium transients in cells where the nuclear membrane permeability was already elevated (Fig.4g). A third group of cells exhibited one or more superphysiologic calcium signals with mostly rounded peaks, and a subsequent increase in nuclear membrane permeability indicating cell death (Fig.4h). In regions greater than 400 µm below the impact site there was a group of cells that exhibited one or more physiologic calcium signals with mostly square peaks, maintained relatively high mitochondrial polarity, and showed no change to the nuclear membrane permeability (Fig.4i).

Finally, we observed two additional groups of cells that maintained low nuclear membrane permeability throughout the experiment. In the first group the mitochondria remained polarized (Fig.4j), while in the second all three signals were low (Fig. 4k). Both of these groups were evenly distributed throughout the impacted samples and the controls.

### II. DECISION TREE METHODOLOGY

### III. TRACKING AND INTENSITY ANALYSIS CODE

This set of codes requires Crocker Grier particle tracking code found here: `https://site.physics.georgetown.edu/matlab/`, and export_fig from the MATLAB Fileshare found here: `https://www.mathworks.com/matlabcentral/fileexchange/23629-export_fig`, as well as the MATLAB Signal Processing Toolbox.

The tracking code does not necessarily re-assign the same cell ID to the same cell each time it is run, due to the nature of the way the Crocker and Grier code is written. Because of this, different sample data is provided for some of the different steps here. This is because the manually-sorted data is not necessarily going to have the same cells with the same ID when the user runs the sample data.

The post-impact input files for this code should be in RGB format .tif files. ImageJ or Fiji can be used to convert from other image formats. Ensure that the colors are in the correct channels (for example, Slidebook likes to swap R and B). The impact input file should be an 8-bit .tif file.

For cells that are not moving, the video can be registered onto the first frame. This will allow the user to connect tracking between the impact and post-impact videos (and only works at the impact site). MATLAB's fitgeotrans and imwarp functions are used to accomplish this. An example snippet of code is provided, but this is not used in the final data analysis so is not integrated into the code as a whole.

To run the codes provided, ensure that the MATLAB directory includes all files within the folder. Start with all_tracking_function_calls. This code will require the user to fill in the various parameters associated with the timing of the images, where the images are saved, alongside the positions that the user wishes to track. Impact and post-impact tracking parameters for the Crocker and Grier algorithm must be filled in, with example values saved within the code itself. Feature extraction parameters should also be filled in (for detailed explanation see the code). Once all parameters are filled, the code can be run. Input parameters for each of the called functions can be found in the functions. Most of the functions require a folder and date to designate where the images are stored and where the output files should be sent to. TrackImpact, TrackPostImpact, and FeatureExtraction have print parameters, which can be set to 'on' or 'off' to determine whether or not the tracking images or feature extraction images are printed. These images are used to help optimize the parameters for tracking or feature extraction. The first time the code is run, the should be set to 'on' in order to ensure that the tracking is working correctly, and that the peak detection looks correct. Future iterations of the code with optimal parameters do not require printing the images. The code takes significantly longer to run if printing all images, but will be faster on a computer with a better GPU (MATLAB renders faster).

Once all_tracking_function_calls is run, the tracking and feature extraction is complete. The user is given a choice between the sorting_function_calls_manual and sorting_function_calls_nomanual codes. For the first time running the data, the user should use the GUI (detailed below) to identify the categories of all of the cells. These categories are then used to label and sort all of the cells. Sorting can be easily accomplished by creating a copy of all intensity curves, then using the first section of sorting_function_calls_manual to make folders corresponding to all of the input categories. ManualDataCompilation will then scrape filenames from within each of these folders to add them to a structure with the correct labels. From here, both of the sorting codes will run CellAttributes to organize features for each time series and DecisionTree to categorize them, finally using SplitPeaksDataCompilation to format the data into shape for time series classification.

The decision tree was custom programmed for our system. After CellAttributes is run in the main script, the features are used to categorize cells into their respective categories. The parameters of the tree must be changed for each new system, with the overall structure remaining quite similar. For other systems, new categories may be identified, and used to determine the if-else statements building the tree.

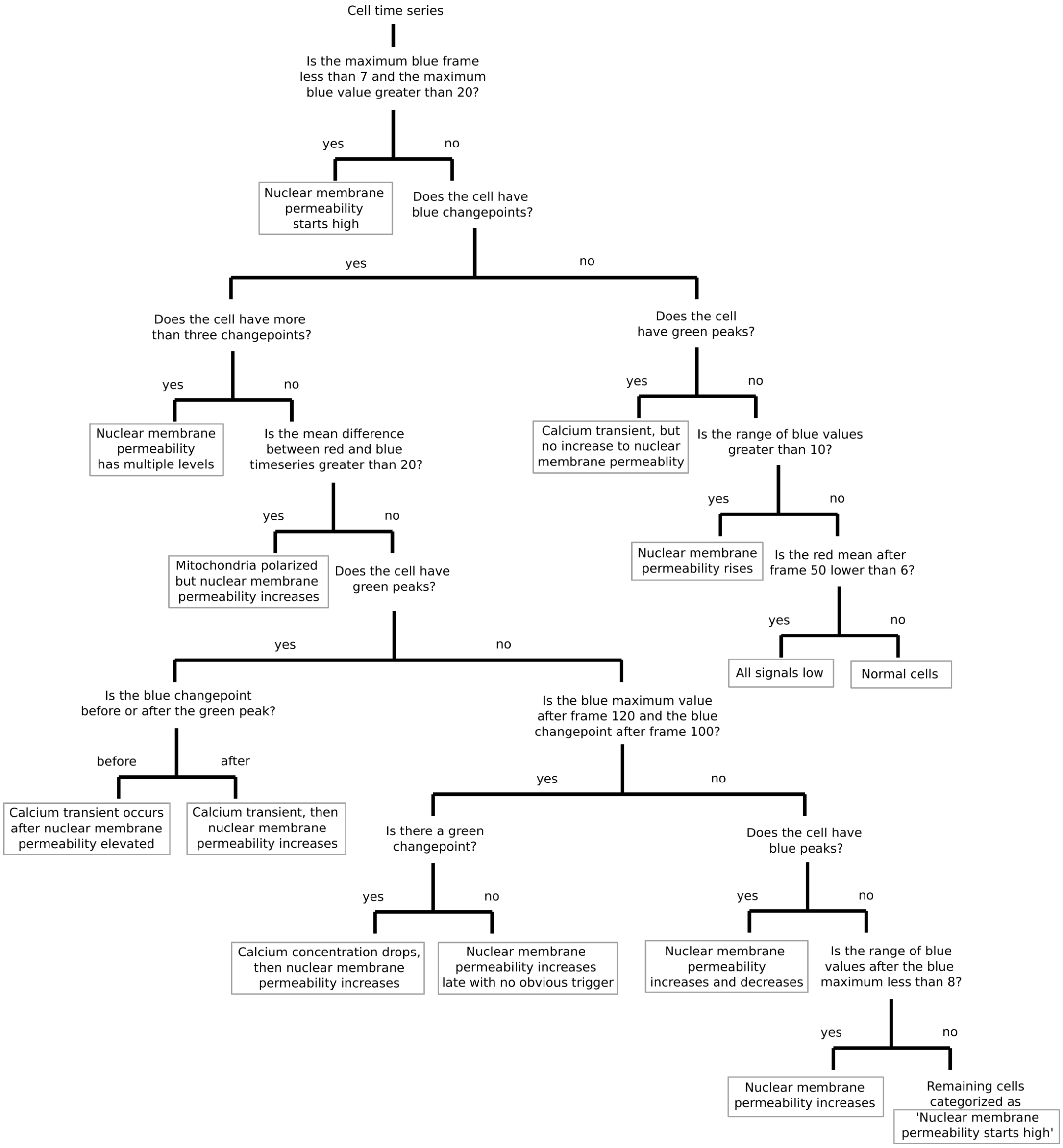


FIG. 1. The full decision tree algorithm for our system. Each cell time series is run through the tree separately. Grey boxes indicate final sorted categories.

### IV. MATLAB GUI

The GUI requires MATLAB 2019a or newer. If other code has been run that has changed figure defaults (like code above), it is recommended that a new instance of MATLAB is opened or the command 'reset(groot);' is run before starting the GUI to avoid figure sizing errors. A video detailing how to use the GUI is included, along with example data.

The GUI has several main functions: to observe cells and groups of cells within the video and plot their individual fluorescence intensities, and to look at how these intensities are distributed. First, pre-processed data is loaded in using the interactive menu and Windows explorer dialog. The pre-processed data consists of the video, tracked intensities of each cell, and location of each cell, with the same naming convention (see example data). Intensity data is taken from 'pos_()_intensity' and 'pos_()_locs' .mat files. Then, the video player in the center can be used to either play at a specified framerate or scroll through all of the frames within the video. Individual cells can be clicked on within the video to display their intensities on the plot side of the GUI. The Cell ID (given by the particle tracking code) or x-y coordinates can be used to find cells as well.

Groups of cells can be selected using the rectangular selection tool. When using this functionality, the plots on the right side of the GUI can be interacted with. First, click within the plot area on one of the three colored plots (but not directly on a line). Then, a cursor will appear.

Click on a point within the graph. After this process is completed, 'highlight cell' can be clicked on to circle the cell within the image, 'Timepoint Histograms' can be clicked on to produce a pop-up with histograms of all three colors at the selected time, and 'Cell Populations' can be clicked on to produce a pop-up with the selected area split into three populations based on the intensity of the blue curve at the selected time. At any point, 'Clear Current Data' can be used to start over. At the end of analysis, 'Save Cell ID List' will pop out a Windows dialog to save the list of cells that were interacted with within this session of the GUI.

### V. TIME SERIES CLASSIFICATION CODE

The hand-sorted data above is used to train several time series classifiers. Data is loaded as a MATLAB .mat file, which is converted to a dataframe. All of the classifier functions are located within classifier_functions.py. In order to train models, labeled data is used, alongside the trainCIF, trainDrCIF, trainROCKET, and trainArsenal functions. These functions will save the models, alongside the accuracy of the model tested on a subset of the data. To use the trained models, the loadCIF, loadDrCIF, loadROCKET, and loadArsenal functions can be used. This will label new data. The model will not be trained exactly the same every time, due to random seeding. However, the accuracies should be very similar.

Dependencies of this code are listed within the functions file. Sktime can be found at `https://www.sktime.org/en/latest/`.

### REFERENCES


1. Choi, J. B. *et al.* Zonal changes in the three-dimensional morphology of the chondron under compression: The relationship among cellular, pericellular, and extracellular deformation in articular cartilage. *Journal of Biomechanics* **40,** 2596–2603. ISSN: 0021-9290 (Jan. 2007).
2. Youn, I., Choi, J. B., Cao, L., Setton, L. A. & Guilak, F. Zonal variations in the three-dimensional morphology of the chondron measured in situ using confocal microscopy. *Osteoarthritis and Cartilage* **14,** 889–897. ISSN: 1063-4584 (Sept. 2006).
3. Guilak, F., Ratcliffe, A. & Mow, V. C. Chondrocyte deformation and local tissue strain in articular cartilage: A confocal microscopy study. en. *Journal of Orthopaedic Research* **13,** 410–421. ISSN: 1554-527X.
4. Welhaven, H. D., McCutchen, C. N. & June, R. K. *A comparison of shear- and compression-induced mechanotransduction in SW1353 chondrocytes* en. preprint (Bioengineering, May 2021).
5. Bartell, L. R. *et al.* Mitoprotective therapy prevents rapid, strain-dependent mitochondrial dysfunction after articular cartilage injury. en. *Journal of Orthopaedic Research* **38,** 1257–1267. ISSN: 1554-527X (2020).
6. O'Conor, C. J., Leddy, H. A., Benefield, H. C., Liedtke, W. B. & Guilak, F. TRPV4-mediated mechanotransduction regulates the metabolic response of chondrocytes to dynamic loading. en. *Proceedings of the National Academy of Sciences* **111,** 1316–1321. ISSN: 0027-8424, 1091-6490 (Jan. 2014).
7. Rocio Servin-Vences, M., Moroni, M., Lewin, G. R. & Poole, K. Direct measurement of TRPV4 and PIEZO1 activity reveals multiple mechanotransduction pathways in chondrocytes. en. *eLife* **6,** e21074. ISSN: 2050-084X (Jan. 2017).
8. Lee, W., Guilak, F. & Liedtke, W. in *Current Topics in Membranes* (ed Gottlieb, P. A.) 263–273 (Academic Press, Jan. 2017).
9. Lee, W. *et al.* Synergy between Piezo1 and Piezo2 channels confers high-strain mechanosensitivity to articular cartilage. en. *Proceedings of the National Academy of Sciences* **111,** E5114–E5122. ISSN: 0027-8424, 1091-6490 (Nov. 2014).
10. Coste, B. *et al.* Piezo1 and Piezo2 Are Essential Components of Distinct Mechanically Activated Cation Channels. en. *Science* **330,** 55–60. ISSN: 0036-8075, 1095-9203 (Oct. 2010).
